# Osa-miR820 regulatory node primes rice plants to tolerate salt stress in an agronomically advantageous manner

**DOI:** 10.1101/2021.01.20.427536

**Authors:** Neha Sharma, Sudhir Kumar, Neeti Sanan-Mishra

## Abstract

Plant microRNAs (miRs) play an important role in regulating gene expression under normal and stressful environments. Here we report the functional implications on the role of Osa-miR820, which can be grouped as a young, rice-specific miR. It is a member of the class II transposon-derived small RNA family and is processed as 21-nt and 24-nt length variants, respectively. Size of the miR820 family varies from 1 to 16 across the Oryza AA genomes. The 21–nt Osa-miR820 negatively regulates a *de novo* methylase, OsDRM2 (*domains rearranged methyl transferase*) that prevents methylation of the CACTA transposon loci in the rice genome. In an earlier report we have detailed the expression profiles of Osa-miR820 and its target in abiotic stress responses using rice varieties exhibiting varying response to salt stress. In this study, artificial miR based approach was employed to specifically overexpress 21-nt Osa-miR820 in rice plants (OX-820). These plants exhibited enhanced vigour, ~25-30% increase in the number of spikelets per panicle and increased grain filling, under normal and salt stress conditions. The OX-820 lines showed a better water use efficiency and higher proline accumulation under salt stress. These plants can serve as a useful source for dissecting the molecular machinery governed by Osa-miR820:DRM2 node to prime tolerance to salt stress in an agronomically advantageous manner.

## 1. Introduction

The sessile nature of plants warrants a continued interaction with their external environment for optimal growth and development. This is a multifarious activity that necessitates intricate coordination of gene expression at the transcriptional and post-transcriptional levels. The microRNAs (miRs) are endogenously transcribed regulatory small RNAs, which act in an exceedingly precise manner to rapidly adjust changes in gene expression. They regulate almost every aspect of plant biology such as root and shoot initiation, leaf development, organogenesis, phase transition, flower morphogenesis, phytohormone response and so on [1–6]. The role of miRs in synchronizing plant responses to biotic and abiotic stresses is well recognized [7–9] and has been extensively studied in model plants, Arabidopsis [5, 7, 9] and rice [8, 10–13]. New insights into the expression profiles and regulation of miRs have resulted from the application of deep-sequencing technologies and large-scale small RNA expression profiling [14–18]. Using the approach of comparative miR profiling followed by experimental validation, our group profiled rice miRs that show tissue-preferential expression patterns [2] and mapped the changes in their profiles in response to salt stress using different rice genotypes [13, 19].

Studies involving miR over-expression also highlighted their crucial roles in regulating gene expression for optimal development and response to stress. It was shown that the expression of miR395 was induced under conditions of sulphate limitation to regulate a high-affinity sulphate transporter and three ATP sulfurylases (ATPS) so that plants could survive in low sulphur containing soils [20]. Overexpression of Osa-miR395h in tobacco impaired sulphate homeostasis and distribution in the transgenic plants [21]. It was also reported that osa-miR156 could negatively control the number of panicle branches and grain yield through its target, OsSPL14 [22]. In addition, pollen-specific miRs along with specific siRNAs were proposed to be key regulators in the phase transition from bicellular pollen to tricellular pollen stage [23]. These observations suggest a role for miRs in controlling plant architecture and yield.

Over-expression of stress regulated miRs revealed their additional functions by generating visible plant phenotypes. The drought responsive Osa-miR393 regulates two auxin receptor genes *Osa-TIR1* and *Osa-AFB2* [24]. Its overexpression resulted in plants that were susceptible to drought and salt stresses [25]. Interestingly these plants showed increased tillering and early flowering [26]. Another example is provided by Osa-miR396d which targets the *growth regulatory factor (grf)* gene to regulate floral organ identity and husk opening [27]. The over expression of miR396 in tobacco confirmed its important roles in drought tolerance and leaf development [28]. Likewise it was seen that overexpression of high-temperature-responsive Osa-miR397 modulates the expression of L-ascorbate oxidase [29, 30], resulting in increased grain size, more rice panicle branching and higher grain productivity [31].

The present study is focused on Osa-miR820 to understand its physiological and functional relevance in rice. It is a young, rice specific miR, that was first reported in undifferentiated embryogenic rice callus tissues as miR583 [32]. Later it was proposed to be a member of the class II transposon derived small RNA family, with its expression being controlled epigenetically at its own locus [33, 34]. Osa-miR820 was shown to be down-regulated under drought [<u>35</u>], salt [<u>36</u>] and arsenic [<u>37</u>] stress. Size of the miR820 family varies from 1 to 16 across the Oryza AA genomes [38] and there are three members in the indica genome [36]. Moreover, 21-nt and 24-nt length variants have also been proposed for Osa-miR820 family in rice, which are processed by the action of the DCL-1 and DCL-3, respectively [39], suggesting its dual mode of action. The 21–nt Osa-miR820 negatively regulates a *de novo* methylase, OsDRM2 (*domains rearranged methyl transferase*) in ‘trans’ to prevent methylation of the CACTA transposon loci in the rice genome [39]. The 24-nt variant act to establish a regulatory loop by controlling epigenetic modifications of OsDRM2 as well as its own locus [<u>34</u>]. These findings indicate a complex regulatory phenomenon governed by the miR family.

Our group has previously described the detailed expression profiles of Osa-miR820 and its target in different tissues of salt-tolerant and salt-susceptible varieties of rice under normal and abiotic stress (salt, high temperature and drought) conditions [36]. The regulatory role of this miR on DRM2 transcripts was evident and the narrow windows of transcript regulation were captured. For gaining further insights into the biological significance of 21-nt species of Osa-miR820 in rice, artificial miR based over-expressing transgenic lines, OX-820 were generated [40]. In the present work, we report the phenotype and physiology of OX-820 under normal and saline conditions.

## 2. Results

### 2.1. Osa-miR820 mapped to QTLs associated with agronomic parameters

To gain insight into the putative role of Osa-miR820, its association with rice QTLs was searched *in silico*. QTL is a large genomic region controlling a specific trait in plants. Earlier studies have correlated the expression of QTL associated miRs with traits related to agronomic performance [41]. Thus, all QTLs overlapping with the three pri-miR820 loci (coding for miR820a, b and c) on rice chromosomes 1, 7 and 10 were identified and mapped (**Supplementary Fig S1**).

This analysis resulted in the identification of 55, 28 and 68 QTLs mapping to Osa-miR820a, Osa-miR820b and Osa-miR820c loci, respectively. The trait association of QTLs with Osa-miR820 loci is depicted in an interaction map shown in **Figure 1A** and the list of Osa-miR820 associated QTLs is provided in **Table 1**. Highest trait density correlated with Osa-miR820b on chromosome 7, followed by Osa-miR820a on chromosome 1 and then with Osa-miR820c on chromosome 10. All three loci of Osa-miR820 showed association with 2 QTLs linked to the trait names *‘Days to heading’* and *‘Plant height’*, respectively. Among the identified QTLs, 11 showed association with more than one trait. Category wise distribution of traits showed that the category *‘Yield’* was maximally represented followed by *‘Vigor’*. This indicates that Osa-miR820 family could be functionally associated with the two growth parameters.

**Figure 1.**
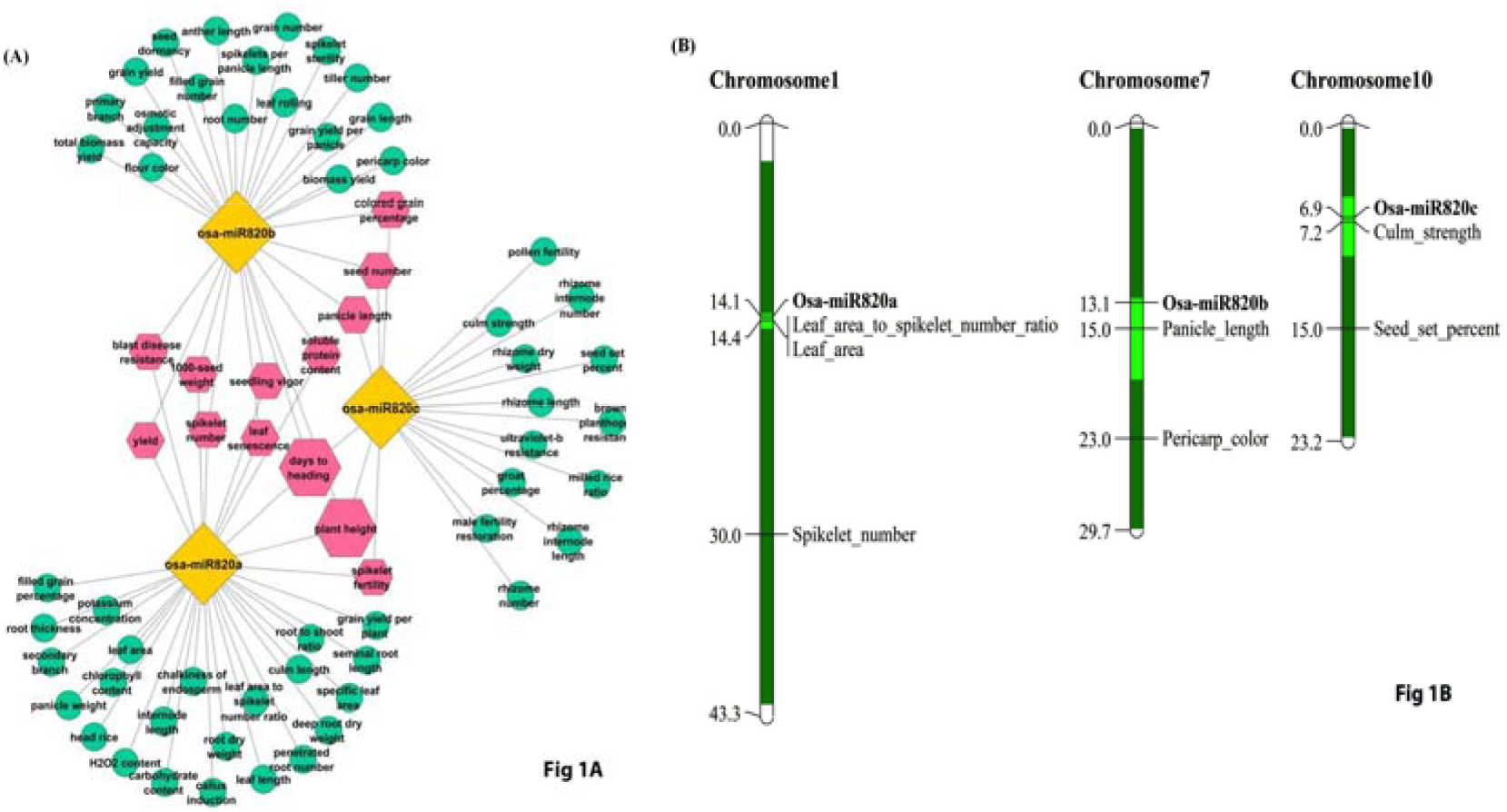
Mapping of QTL associated traits to miR820 locus. A) Cytoscape based interaction map. Yellow boxes denote the miR family member; green boxes denote the trait for the associated QTL; Pink hexagons denote QTLs associated with more than one miR820 family member. B) Karyoview of largest and smallest QTLs identified for Osa-miR820a, b and c.

**Table 1.** Trait wise listing of number of QTLs associated with Osa-miR820 locus.

| Trait category | Chromosome 1<br>(Osa-miR820a) | Chromosome 7<br>(Osa-miR820b) | Chromosome 10<br>(Osa-miR820c) |
| --- | --- | --- | --- |
| Yield | 19 | 18 | 5 |
| Vigor | 8 | 18 | 2 |
| Development | 6 | 13 | 2 |
| Anatomy | 8 | 6 | 6 |
| Quality | 2 | 8 | 4 |
| Biochemical | 7 | 1 | 0 |
| Abiotic stress | 3 | 2 | 2 |
| Sterility or fertility | 1 | 1 | 4 |
| Biotic stress | 1 | 1 | 3 |
| **Trait for smallest QTL** | Ratio of leaf area to<br>spikelet number | Panicle length | Culm length |

To be more specific in predicting the traits associated with Osa-miR820 function, the smallest QTL on each locus was identified (**Fig. 1B**). This analysis identified *“ratio of leaf area to spikelet number”, “panicle length”* and *“culm length”* as the associated traits (**Table 1**). These traits define agronomic parameters that can directly or indirectly affect the rice crop yield. Thus, it was speculated that Osa-miR820 might have a role in regulating plant biomass and/or productivity.

### 2.1. OX-820 rice plants exhibit increased tillering, panicle vigour and enhanced grain yield

The presence of two length variants of Osa-miR820 of 21-nt and 24-nt, respectively having common targets presented an interesting challenge in understanding the functional significance of this miR in the biology of the rice plant. Therefore, artificial miR based strategy was adopted to specifically over-express the 21-nt length variants. The Osa-miR820 overexpressing transgenic lines (OX-820) were generated and confirmed, as described earlier [40].

For phenotypic comparisons, healthy mature seeds of OX-820 and wild type (WT) were grown on germinating sheets in greenhouse conditions. It was observed that OX-820 seedlings were relatively longer and healthier as compared to WT (**Fig. 2A**). The increase in length was also evident on comparing the roots (**Fig. 2A**). The increase in OX-820 plant height was observed in the mature plants as well (**Fig. 2B**), but the plants exhibited a decrease in total biomass dry weight (**Table 2**). On comparing the leaf characteristics, it emerged that the leaf length was more in OX-820 plants as compared to the WT but there was no change in the leaf width (**Table 2**). The OX-820 rice plants showed stouter roots with overall decrease in root length but slightly increased root dry weight (**Table 2**).

**Figure 2.**
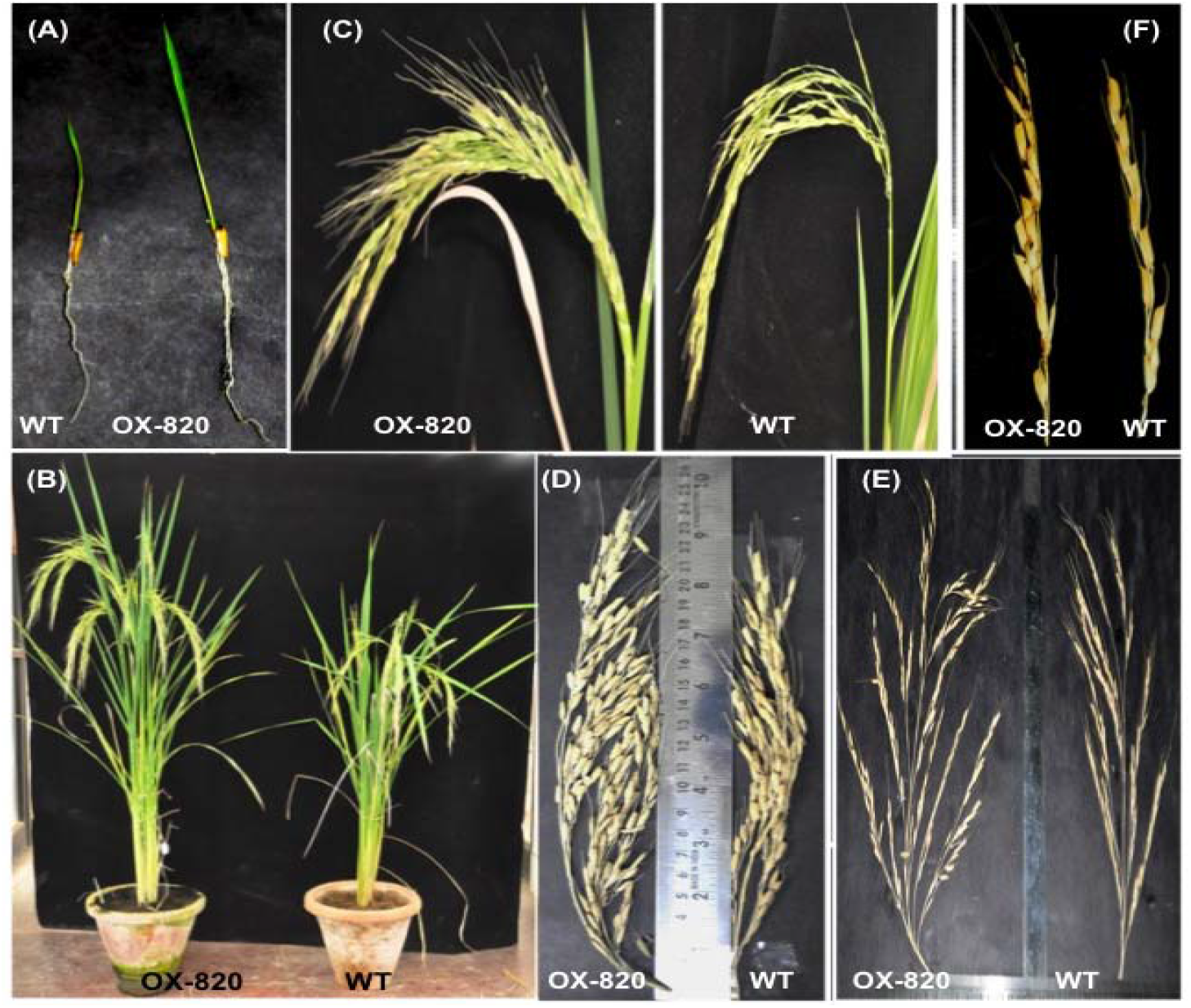
Phenotype of OX-820 rice plants as compared with wild type (WT) (A) 10-day old rice seedlings (B) Plants at the maturity stage (C) Panicles at early grain filling stage (D) Panicle length (E) Panicle branching and (F) Grain filling in an individual spikelet.

**Table 2.** Comparison of vegetative growth related traits in OX-820 transgenic and WT plants.

| Line | Primary tillers<br>(number) <sup>#</sup> | Leaf<br>length<br>(cm) <sup>##</sup> | Mature<br>plant height<br>(cm) <sup>##</sup> | Plant biomass<br>dry weight<br>(g) <sup>##</sup> | Mature<br>root length<br>(cm) <sup>##</sup> | Mature root<br>dry weight<br>(g) <sup>##</sup> |
| --- | --- | --- | --- | --- | --- | --- |
| WT-1 | 6 | 45.0±1.3 | 34.5±1.4 | 26.0±0.5 | 25.5±1.3 | 4.38±0.1 |
| WT-2 | 5 | 42.8±1.2 | 33.7±1.3 | 25.4±0.5 | 23.6±1.2 | 4.0±0.1 |
| WT-3 | 5 | 44.2±1.3 | 31.9±1.3 | 25.8±0.5 | 24.7±1.2 | 4.25±0.1 |
| OX-820-L1 | 8 | 59.5±1.8 | 37.4±1.5 | 21.5±0.4 | 22.2±1.1 | 4.7±0.1 |
| OX-820-L2 | 7 | 56.7±1.7 | 39.6±1.6 | 18.0±0.6 | 25.4±0.9 | 5.02±0.1 |
| OX-820-L3 | 8 | 56.0±1.7 | 38.8±1.5 | 23.5±0.5 | 21.0±1.3 | 4.8±0.1 |
| OX-820-L4 | 8 | 54.0±1.6 | 38.3±1.5 | 22.2±0.4 | 19.5±1.0 | 4.28±0.1 |

At maturity the OX-820 rice plants had more number of primary tillers (**Fig. 2B**) and robust panicles (**Fig. 2C**), when assessed with respect to the WT. The increase in number of tillers was in direct correlation with the number of panicles (**Table 3**). Analysis of individual panicles revealed that in OX-820 panicles were longer (**Fig. 2D**), profusely branched on their primary and secondary axis (**Fig. 2E**) and exhibited denser grain filling (**Fig. 2F**). This resulted in increased drooping of panicles at the post-anthesis stage in OX-820 plants (**Fig. 2C**). Thus, it was evident that constitutive over-expression of the 21-nt sequence of Osa-miR820 generated a distinct phenotype in rice indicating its role in modulating plant architecture.

**Table 3.** Comparison of agronomic traits in OX-820 transgenic and WT plants.

| Line<br>number | No. of<br>panicles<sup>#</sup> | 1°branches<br>per<br>panicle<sup>#</sup> | Panicle<br>length<br>(cm)<sup>#\$</sup> | No. of<br>spikelets per<br>panicle<sup>##</sup> | Total no.<br>of grains<sup>#</sup> | 1000-seed<br>weight<br>(g)<sup>##</sup> | Grain<br>length<br>(cm)<sup>##</sup> | Grain<br>width<br>(cm)<sup>##</sup> |
| --- | --- | --- | --- | --- | --- | --- | --- | --- |
| WT-1 | 6 | 6 | 24.1±0.5 | 136±6.8 | 780±31 | 20.9±0.8 | 10.0 | 2.1 |
| WT-2 | 5 | 5 | 24.9±0.5 | 131±6.6 | 647±26 | 19.8±0.8 | 10.1 | 2.2 |
| WT-3 | 6 | 6 | 23.7±0.5 | 128±7.2 | 804±32 | 20.2±0.8 | 10.9 | 2.2 |
| OX-820-L1 | 9 | 5 | 28.2±0.5 | 159±8.0 | 1104±44 | 23.4±0.9 | 11.0 | 2.1 |
| OX-820-L2 | 8 | 8 | 29.5±0.6 | 168±8.4 | 1206±48 | 23.2±0.9 | 11.65 | 2.4 |
| OX-820-L3 | 9 | 7 | 26.4±0.5 | 171±8.5 | 1373±55 | 22.6±0.9 | 10.8 | 2.3 |

High tillering capacity is an important agronomic trait in cereals and it has a positive correlation with grain yield in rice. However higher number of tillers tends to create an imbalance of photosynthetic assimilates among vegetative and reproductive parts, which may compromise productivity [42, 43]. The OX-820 transgenic plants were compared for agronomic parameters to evaluate their grain productivity (**Table 3**). It was observed that each primary tiller contained a dense panicle with higher number of spikelets as compared to WT. There was 25-30% increase in the number of spikelets per panicle in OX-820 plants. This increase was largely due to enhancement in the length and branching of panicles. The spikelets showed increased grain filling resulting in 50-60% increase in the number of grains (**Table 3**). The individual grain width remained unaffected and there was a marginal increase in grain length. Thus, constitutive over-expression of the 21-nt sequence of Osa-miR820 resulted in increase in number of spikelets and improvement in grain filling per panicle leading to enhanced grain yield.

### 2.2. Overexpression of Osa-miR820 does not influence response to salt stress

Earlier studies have shown that Osa-miR820 levels were down regulated under salt stress [<u>36</u>], so it was apparent to evaluate the effect of salt stress on OX-820 plant phenotype and physiology. To understand the response of OX-820 plants to salt stress the chlorophyll retention assay [<u>44</u>], was performed. Discs cut from healthy leaves of both OX-820 and WT plants were kept in different concentrations of NaCl and monitored at regular time intervals. Discs kept in deionized water served as control. After 72 h, the leaf discs were collected (**Fig. 3A**) and the chlorophyll content was quantitated and plotted (**Fig. 3B**). Significant difference was not observed between leaf discs of OX-820 and WT plants, indicating that over-expression of Osa-miR820 does not impart tolerance or enhance sensitivity of the plants to salt stress.

**Figure 3.**
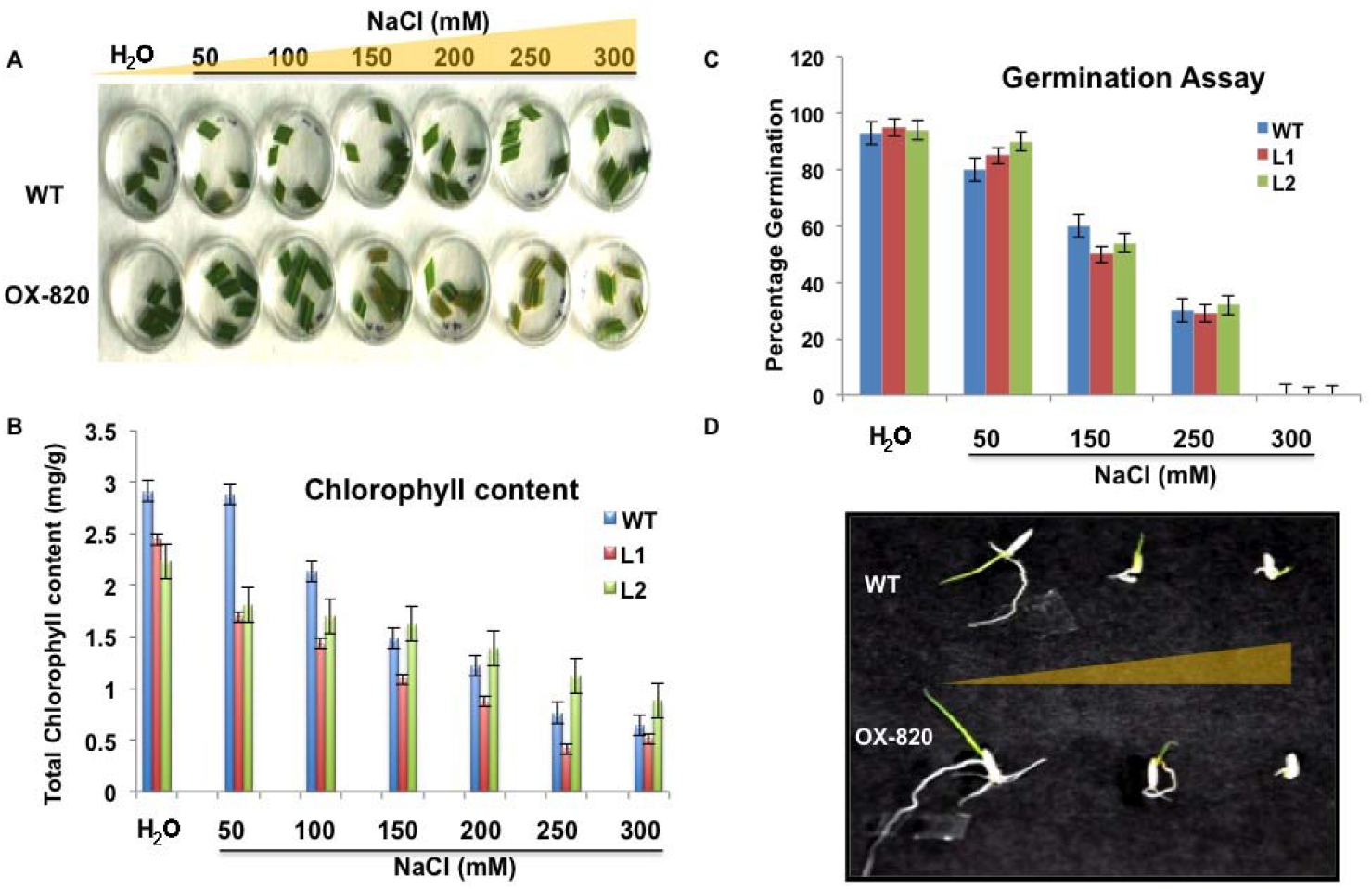
Assaying the effect of salt stress on OX-820 and WT plants. **(A)** Representative picture of leaf discs placed in increasing concentration of NaCl solution to show chlorophyll retention. (B) Plot of chlorophyll content estimated from leaf discs after 72 h of stress (C) Percentage germination under increasing salt gradient (D) Representative picture showing phenotype of germinating OX-820 and WT seeds in presence of water, 150 mM NaCl and 250 mM NaCl.

#### 2.2.1. Assay for germination and seedling growth

To follow the effect of salt stress on germination, seeds of WT plants and OX-820 lines were germinated in presence of different NaCl concentrations and the percentage values were plotted (**Fig. 3C**). Seed germination was negatively impacted in presence of salt stress and efficiency of germination decreased by 35-40% in presence of 250 mM NaCl (**Fig. 3D**). At 300 mM NaCl, both OX-820 and WT seeds failed to germinate. This indicated that over-expression of Osa-miR820 did not influence their sensitivity towards salt.

The growth performance of OX-820 seedlings under salt stress was also analysed, by measuring the total fresh weight (**Fig. 4A**), shoot length (**Fig. 4B**) and root length (**Fig. 4C**) of seedlings stressed with NaCl for 1 week. Under unstressed conditions (water grown) both the WT and transgenic seedlings grew well. The OX-820 seedlings were longer than WT and accumulated higher fresh weight. In the presence of NaCl the inhibition in growth of OX-820 seedlings was similar to the levels observed in WT. Seedling growth was severely retarded at NaCl concentrations higher than 200 mM. Thus, OX-820 seedlings like their WT counterparts could tolerate mild salinity conditions, but could not resist high salt concentrations.

**Figure 4.**
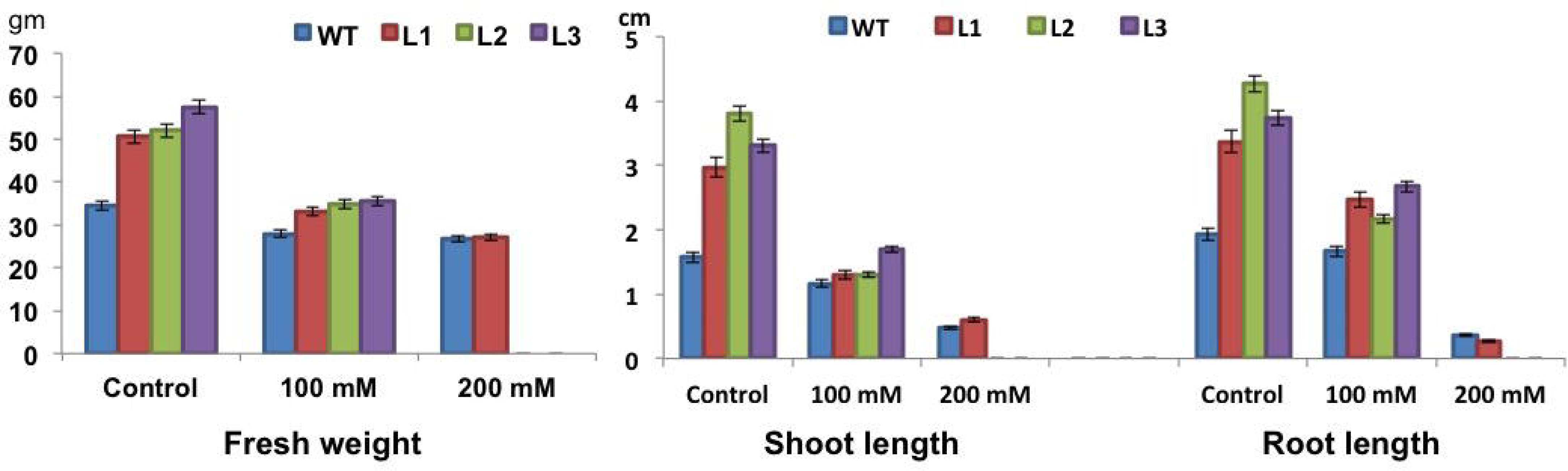
Effect of salt stress on OX-820 and WT seedlings. The graphs show measurements of (A) fresh weight (B) shoot length and (C) root length. The unit of measurement of weight is grams and unit of measurement of length is centimetre.

#### 2.2.2. Salt stressed OX-820 plants resemble un-stressed WT plants

To understand the response of plants to long term salt stress, the OX-820 and WT plants were grown till maturity by supplying 250 mM NaCl solution once every 15 days. The plants grown in normal watered pots served as controls. The plants were assessed for various morphological, physiological and grain yield parameters.

Under control conditions, OX-820 plants were taller with more number of tillers as compared to WT. Under salt stressed conditions, the OX-820 plants showed marked reduction in plant height, but there was no significant change in the number of tillers (**Fig. 5A**). In contrast the salt stressed WT plants did not show reduction in plant height but the number of primary tillers were drastically reduced. On analysing the phenotype of mature roots it was observed that under control conditions, OX-820 plants had profusely branched roots and enhanced root biomass as compared to WT plants (**Fig. 5B**). Exposure to salt stress reduced the root biomass in both WT and OX-820 plants.

**Figure 5.**
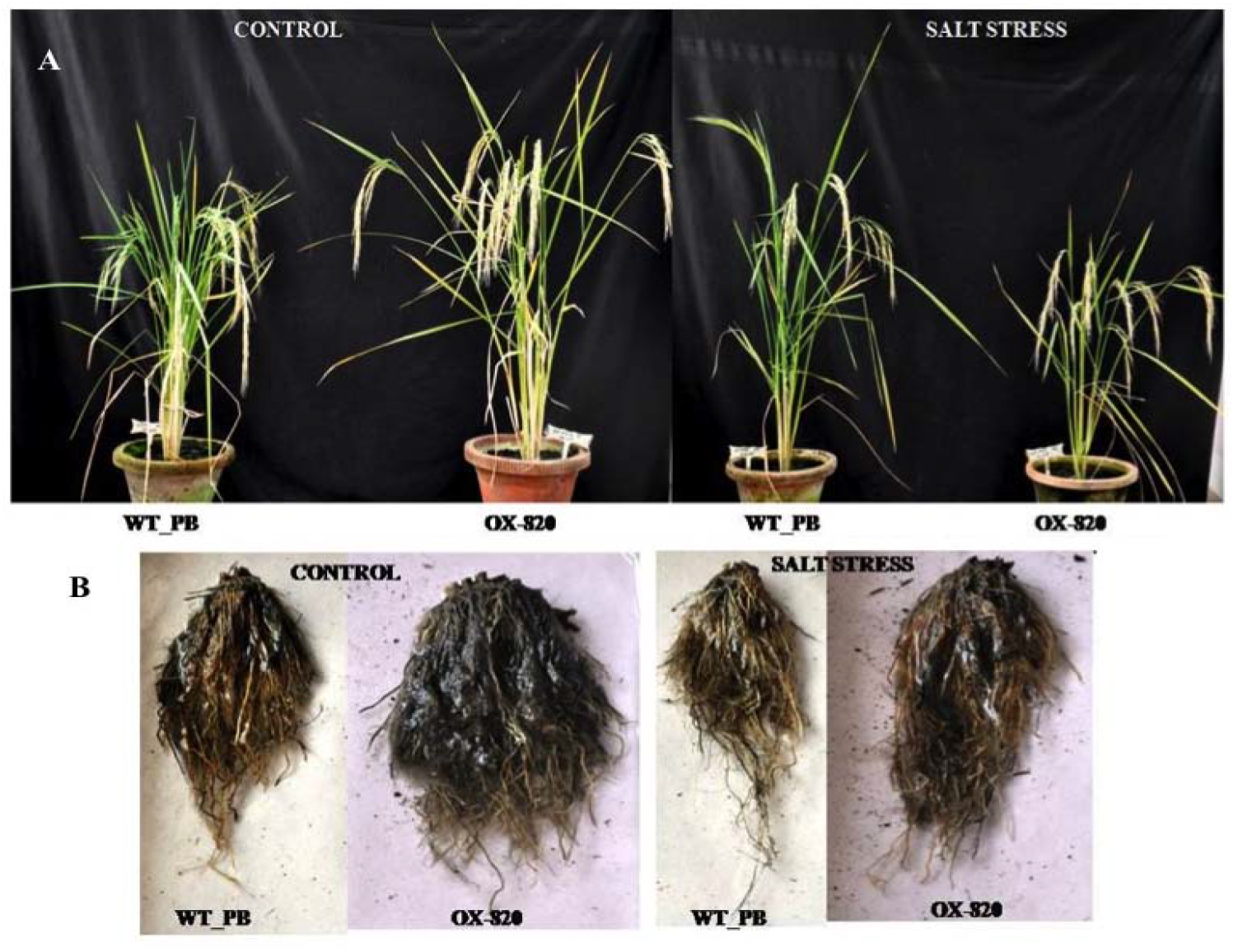
Phenotype of OX-820 and WT plants under long term salt stress. (A) mature plants (B) roots NaCl.

The OX-820 plants were negatively affected by salt stress, but the extent of reduction was much less than that observed in salt stressed WT plants. Interestingly the overall salt stressed phenotype of OX-820 plants appeared similar to that of unstressed WT plants (**Fig. 5**). This is also apparent from the measurements for plant height (**Fig. 6A**), leaf length (**Fig. 6B**), biomass dry weight (**Fig. 6C**), primary tiller numbers (**Fig. 6D**), total number of panicles per plant (**Fig. 6E**) and total panicle weight per plant (**Fig. 6F**).

**Figure 6.**
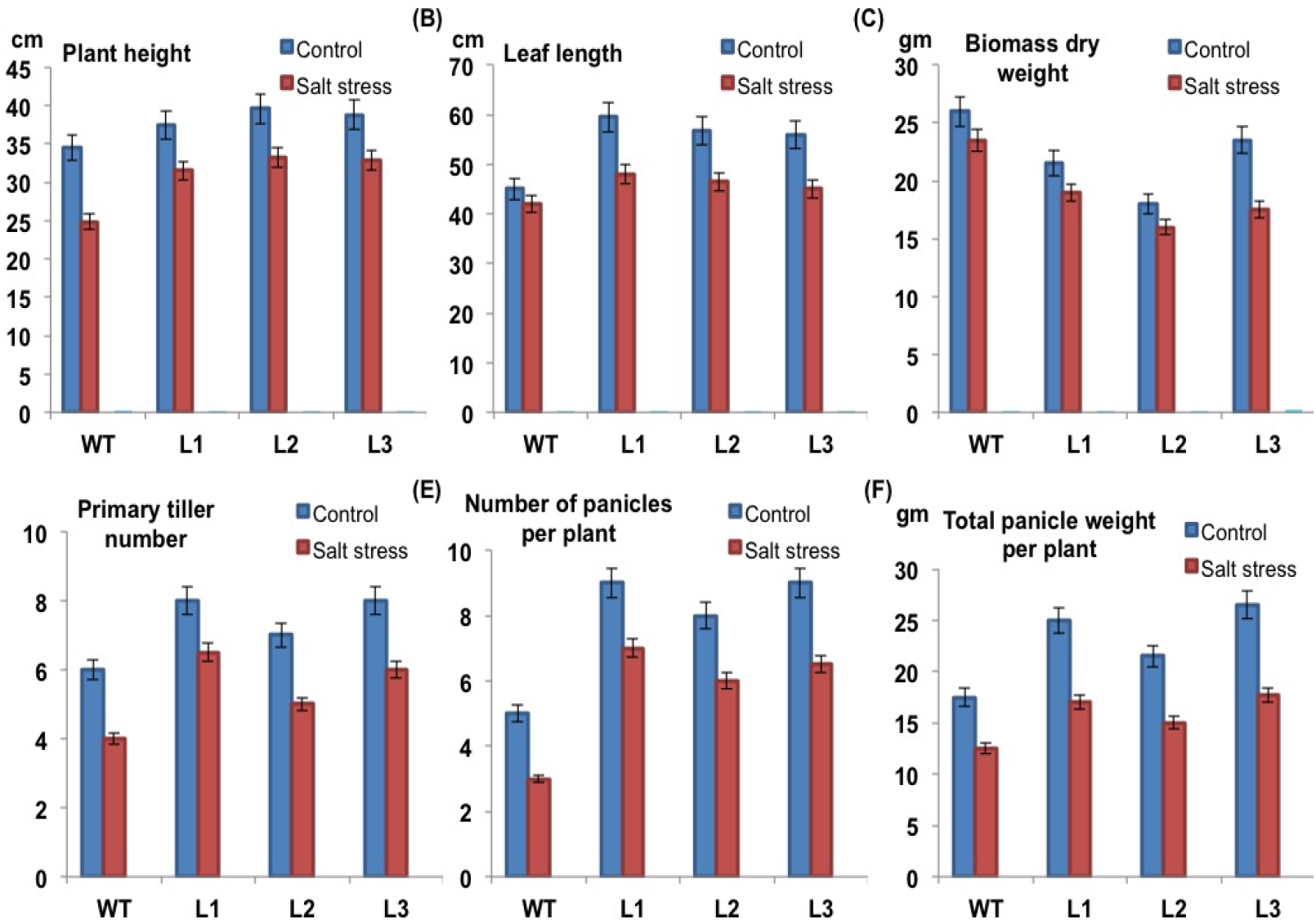
Comparative analysis of vegetative growth and grain yield parameters of mature OX-820 and WT plants grown under normal and salt stress conditions. **(A)** plant height **(B)** leaf length **(C)** biomass dry weight **(D)** primary tiller number **(E)** number of panicles per plant and **(F)** total panicle weight per plant.

### 2.3. OX-820 plants are better adapted to long term salt stress

To understand the effect of salt stress on physiology of OX-820 plants their photosynthetic efficiency and gas exchange measurements were recorded and analysed. Infra Red Gas Analysis (IRGA) of CO_2_ is most widely used, reliable, accurate and simple method for measuring gas exchange and related characteristics. Plant performance was analysed by calculating photochemical efficiency (Fv/Fm) of PSII (**Fig. 7A**), quantum yield (**Fig. 7B**), stomatal conductance (**Fig. 7C**) and transpiration rate (**Fig. 7D**). Quantum yield is the actual efficiency of light utilization during photosynthesis, representing the number of moles of CO_2_ fixed per mole quantum of light absorbed by a leaf [45].

**Figure 7.**
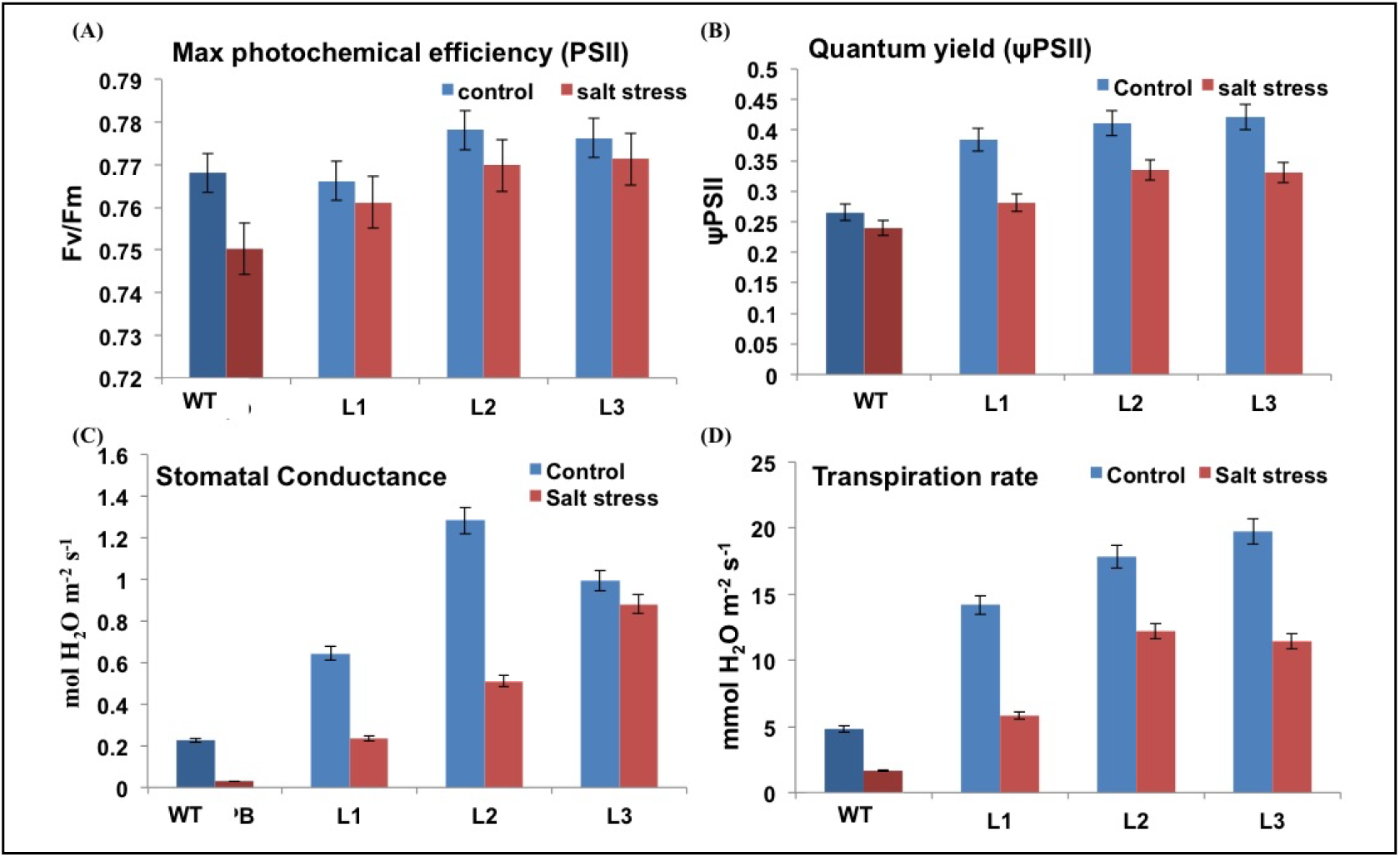
Comparative analysis of photosynthetic and gaseous exchange parameters of mature OX-820 and WT plants. **(A)** Fv/Fm **(B)** quantum yield **(C)** Stomatal conductance **(D)** transpiration rate.

The Fv/Fm ratios of OX-820 and WT leaves were similar under control and salt stress conditions (difference of 0-0.5 is considered negligible). The imposition of stress caused a slight decrease in Fv/Fm ratios in both cases. The quantum yield, stomatal conductance and transpiration rates for OX-820 plants were higher as compared to the WT under unstressed conditions. In salt stressed OX-820 plants these values declined and became comparable to those of unstressed WT plants (**Fig. 7B-D**). The transgenic lines showed better photosynthetic and water use efficiency over the WT, which indicates that OX-820 were better adapted to retain water during salt stress. This could be one of the reasons behind their improved performance under long term salt stress.

Free proline measurement is a well studied parameter for measuring the salt stress tolerance capacity in plants [46]. The OX-820 plants exhibited higher proline levels as compared to WT under unstressed conditions (**Fig. 8**). Exposure to salt stress caused a sharp decline in proline content, but the levels were relatively higher in OX-820 plants. Higher free proline content in salt stressed OX-820 transgenics indicates the activation of a protective cellular machinery that helps the plants to withstand the toxic effect of salt.

**Figure 8.**
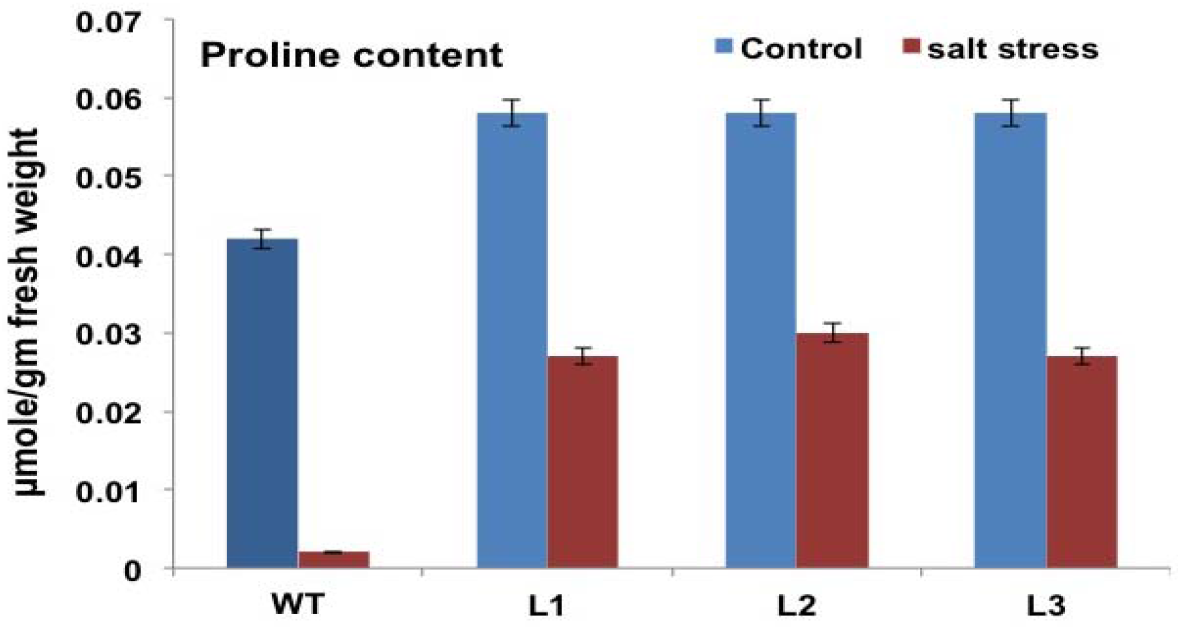
Measurement of free proline content in OX-820 and WT plants.

## 3. Discussion

Cataloguing of miRs in rice is an important step towards understanding the physiology of this important cereal crop. A large number of miRs have been discovered in rice through experimental and computational approaches but functional significance of majority of them is still unknown. Osa-miR820 constitutes a young, rice specific miR family and its three closely related members, Osa-miR820a (MI0005263), Osa-miR820b (MI0005264) and Osa-miR820c (MI0005265) are most conserved in all *Oryza* AA genomes. In addition, at least 52 novel members of this family have been identified across *Oryza* AA genomes and 11 from *Oryza punctate* [38]. Size of the miR820 family varies from 1 to 16 across the Oryza AA genomes [38] and there are three members in the indica genome [36]. These three loci generate both 21-nt and 24-nt length variants by the action of the DCL-1 and DCL-3, respectively [39], suggesting the operation of complex regulatory loops by the miR family.

To gain rapid indications on the functions of this miR, the three pri-miR820(a,b,c) loci were integrated with known rice QTLs. Among the co-mapped QTLs, two QTLs associated with the traits *‘Days to heading’* and *‘Plant height’* showed interaction with all the three loci. The data obtained indicated that prospective roles of this miR family might be related with regulation of plant vigour and yield. Artificial miR based strategy was adopted to constitutively over-express the 21-nt length variants of Osa-miR820 [40]. This generated a distinct phenotype as OX-820 seedlings and mature plants were relatively longer and healthier as compared to WT. Each OX-820 rice plant had 7-8 primary tillers and robust panicles. Individual OX-820 panicles were longer, profusely branched on their primary and secondary axis and exhibited 25-30% more spikelets per panicle. It can be speculated that the robust morphology of OX-820 roots may be providing improved regulation of nutrient and water absorption from the soil for its better survival. The study of primary grain yield components revealed that in OX-820 plants the panicle weight was augmented due to 50-60% increase in grain filling. Thus, OX-820 plants exhibited improved grain filling and enhancement in total grain yield. These observations indicated a crucial role for this miR in modulating plant architecture.

The enhanced vigor and higher grain yield attained by OX-820 plants could broadly explain the association of Osa-miR820 loci with QTLs for traits such as panicle length, culm length and leaf area to spikelet ratio. In general, high yielding varieties have a higher spikelet to leaf ratio indicating higher source to sink capacity for such cultivars [43,47]. It was evident that overexpression of Osa-miR820 increased the level of branching in plants which served as a major determinant for the enhanced tillers, root biomass and panicle vigour. Morphogenetic processes in rice plants are sequential and highly synchronized such that tiller buds arise at the apical end after the formation of flag leaf. The profuse branching pattern of OX-820 plants indicates that Osa-miR820 may be acting upstream in the regulatory cascades.

The miRs not only regulate plant growth and development but also enable the plants to cope up with stressful environment. Soil salinity is one of the most damaging stresses that negatively impacts overall growth and performance of rice, though varietal differences may be observed [48–50]. Salt susceptible cultivars usually show reduced biomass and low crop yields whereas salt tolerant ones may thrive better with less yield penalties, during salinity stress. Earlier studies on salt susceptible Pusa Basmati 1 and salt tolerant Pokkali rice plants had showed that Osa-miR820 was down-regulated under salt and drought stress [<u>36</u>]. Assays for leaf disc chlorophyll retention, germination and seedling growth indicated that over-expression of Osa-miR820 does not impart tolerance or enhance sensitivity of the plants to salt stress. The OX-820 plants, like their WT counterparts, could tolerate mild salinity conditions, but could not resist high salt concentrations.

Analysis of various vegetative and agronomic parameters revealed that the salt stressed OX-820 plants appeared similar to those of unstressed WT plants. Under salt stress, the transgenic lines displayed better photosynthetic performance as indicated by higher values of photochemical efficiency (PSII), and quantum yield. The plants also had improved water relations as indicated by the measurement of stomatal conductance and transpiration rate. Salt stress creates a physiological water deficit leading to stomatal closure thereby affecting the net transpiration rates per unit area of leaf [51,52], therefore stomatal conductance is inversely proportional to stress. The OX-820 plants also exhibited higher proline levels as compared to WT. Proline is a well known osmo-protectant that accumulates in plants during stress and aids in maintenance of membrane integrity, protein stability and cellular homeostasis during adverse environments [53,54]. These observations indicate that Osa-miR820 regulated nodes can prime tolerance to salt stress in an agronomically advantageous manner.

The Osa-miR820 negatively regulates a DNA cytosine methyltransferase, OsDRM2 (*domains rearranged methyltransferase 2*) to prevent *de novo* methylation of C-5 residue of cytosine in the rice genome [39]. This plays a major role in curtailing the transposons and epigenetic activity during reproductive development and stress [55,56]. EST analysis also identified a putative NAC domain containing protein 77 (N77) and a putative aquaporin, plasma membrane intrinsic protein 1-5 (PIP1-5), as targets of Osa-miR820 [56]. The Arabidopsis homolog of DRM2 (uniprot id: Q9M548) has been well characterized. On searching the putative interacting partners of AtDRM2 using ‘IntAct’ [57], a database repository containing information on protein interactions, eight interacting proteins were identified. These included AGO4, AGO9, MSI4 (WD-40 repeat containing protein), HDT3 (histone deacetylase 3), TRP2 (topless related protein 2), SKP1A (SKP1-like protein 1A), H2B6 (Histone H2B.6) and ZOP1 (Zinc Finger Protein 1) protein. MSI4 is a core histone binding protein that positively regulates flowering time by promoting transcriptional repression of FLC, the flowering repressor gene [58]. HDT3 is a histone deacetylation protein whose over-expression leads to increased salt and drought tolerance in Arabidopsis [59]. TRP2 is a member of the protein family that modulates gene expression in multiple processes such as stress response, hormone signalling including auxin and jasmonic acid signalling, as well as flowering [60,61]. It may play a role in meristem specification in plants. Another important interacting partner the SKP1-A is a subunit of E3 ligase and a part of the SCF complex that is involved in ubiquitination and subsequent degradation of proteins. It indirectly influences diverse biological processes including auxin signalling pathway, vegetative and floral organ development, light signalling, embryogenesis and post embryonic development [62–64]. These reports suggest that Osa-miR820 might be indirectly manipulating important cellular mechanisms in rice through its target OsDRM2.

## 4. Conclusion

It can be concluded that over-expression of 21-nt Osa-miR820 in rice increased the plant length and level of branching, which served as major factors for their improved overall productivity by boosting tillering and augmenting panicle vigour. The OX-820 plants exhibited improved photosynthetic performance, water use efficiency and proline content. Though this did not substantially improve the tolerance of plants to salt stress, but in comparison to WT plants the overall plant biomass and tillers were relatively less affected. This manifested in a higher net panicle weight leading to a lowering of yield penalty. Therefore, the complex genetic loops regulated by the miR family modulate plant architecture and prime tolerance to salt stress in an agronomically advantageous manner. The molecular understanding of the underlying mechanisms needs to be further studied for the functional dissection of roles played by Osa-miR820.

## 4. Materials and Methods

### 4.1. Plant Material and stress treatment

Rice plants were grown under controlled conditions at 28±2°C temperature, 70% relative air humidity and 16/8-h light/dark cycle. Seeds were collected from the WT (control) and OX-820 (artificial miR820 overexpressing)) T0 and T1 transgenics. For each analysis, 10-15 mature dehusked rice seeds of each line were surface sterilized and used. NaCl solution for specified concentrations were used to provide salt stress. For providing long term salt stress, the plants were supplied with 250 mM NaCl solution once every 15 days till maturity. The plants grown in normal watered pots served as controls.

### 4.2. Measurement of growth and yield related parameters

At maturity, rice plants were harvested and various grain yield parameters were measured. In each set 6 independent transgenic lines (T0 and T1) were used to record each parameter and standard deviation was calculated for each of them. 10-15 transgenic (PCR confirmed) seedlings from each of these lines were screened and grown. For each line, six plants were subjected to similar recordings under control and salt stressed conditions. The p–test values results shows the significant difference between WT and transgenic lines.

### 4.3. QTL mapping

A list of QTLs mapping to rice chromosomes 1, 7 and 10 were downloaded from Gramene database [65] and these were re-mapped to the rice chromosome using rice genome TIGR ver 7. The overlapping co-ordinates between the miR820 locus and the downloaded sequences led to the identification of putative QTLs associated with Osa-miR820 loci. Based on co-ordinate overlaps, the largest and smallest QTLs on each chromosome were also identified. The workflow for QTL mapping is described in Supplementary Fig S1. A visual interaction map was plotted for all the miR820 associated QTLs using Cytoscape 2.8.3 [66].

### 4.4. Germination and seedling growth

The assay was performed with seeds obtained from T1 plants. For this, 2 sheets of filter paper soaked in water or NaCl solution were placed on the petri dish and sterilized seeds were kept on it. The filter papers were replaced at an interval of two days to prevent salt accumulation. After 10 days, the number of seeds showing the emergence of plumule and radical were recorded. The germinated OX-820 seeds were confirmed for the presence of amiRNA construct. Percentage germination of was calculated by the following formula (Equation 2):

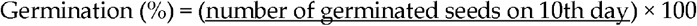

The root and shoot length of 5-10 randomly selected seedlings were measured at 7th day after germination. Fresh weight of the same seedlings was also measured with the help of weighing balance. The OX-820 seedlings were confirmed for the presence of amiRNA construct. All the experiments were done in duplicates and repeated thrice to ensure reproducibility.

### 4.5. Leaf disc assay and Chlorophyll estimation

For this assay 6th leaf of the main culm was used to maintain uniformity. The leaf was cut into pieces and these were placed on petri dishes containing NaCl solution of specified concentrations. Water was used as control. The leaves were floated on the solution with their abaxial surface downwards. Leaf pieces were collected at different time intervals and their chlorophyll content was measured. For each line, six independent biological replicates were taken for different salt stress and control conditions

Extraction of chlorophyll from the leaf tissues was done as per the protocol [44]. 1 g leaf tissue was homogenized in liquid nitrogen and 5 ml of 80% acetone was added to it. It was allowed to thaw at room temperature and then centrifuged at 5000 rpm for 5 min, at room temperature. The green coloured supernatant was collected in a fresh tube. Absorbance was recorded at both 663 nm (A663) and 645 nm (A645), using 80% acetone as a blank. The chlorophyll content was expressed in mg per gram fresh weight using the following formula (Equation 1):

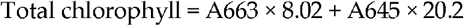

### 4.6. Measurement of gas exchange and photosynthetic parameters

Infra-red gas analyzer (IRGA), a hand held portable, open flow gas exchange instrument was used to measure various gas exchange parameters between the leaf sample and the measuring air chamber. The Net photosynthetic rate (PN) and stomatal conductance (gs), and other associated photosynthetic parameters were recorded in fully expanded leaves using an infrared gas analyzer (IRGA, LiCor, Lincoln, NE) on a sunny day between 10:00 and 11:00. The atmospheric conditions during the measurement were photo synthetically active radiation (PAR), 1,000 ± 5 μmol m−2 s−1, relative humidity 66 ± 5%, atmospheric temperature 25 ± 2°C and atmospheric CO2, 355 μmol mol−1. All the gas exchange parameters were calculated by the LI-6400’s software ver. 6.0. For each line, six independent biological replicates were taken for both salt stress and control conditions.

### 4.7. Proline estimation

Proline was determined using the method described earlier [67]. Briefly, fresh leaf tissues were homogenized in 10 ml sulphosalicyclic acid in ice cold bath, using a pestle and mortar. Homogenate was centrifuged at 10,000 g for 15 min. Then 2 ml filtrate was mixed with 2 ml acid ninhydrin and 2 ml glacial acetic acid. The mixture was incubated at 100°C for 1 h until the coloured complex is developed in water bath. The reaction was terminated by rapid cooling on ice. Then 4 ml toluene was added to the coloured complex and reaction mixture was mixed by vortexing for 15-20 s. Optical density of layer with chromophore (red-pink in colour) was read at 520 nm. Proline content was estimated by using L-Proline standard curve.

## Supplementary Materials

**Supplementary Figure 1** The workflow for QTL mapping.

## Authors’ contributions

Conceptualization, N.S.-M.; Methodology, N.S.; Software, N.S.; Validation, N.S. and S.K.; Formal Analysis, N.S. and S.K.; Investigation, N.S. and S.K.; Resources, N.S.-M.; Writing – Original Draft Preparation, N.S.; Writing – Review & Editing, N.S.-M.; Supervision, N.S.-M.; Project Administration, N.S.-M.; Funding Acquisition, N.S.-M.

## Funding

NSM is thankful for the funds received from International Centre for Genetic Engineering and Biotechnology (ICGEB), New Delhi and Department of Biotechnology (DBT), Government of India.

All authors have approved the final manuscript for publication.

## Data Availability Statement

The data presented in this study are available on request from the corresponding author.

## Acknowledgements

Authors thank Mr. Sandeep Panchal for help with raising the transgenics. NS and SK are thankful to Council of Scientific and Industrial Research (CSIR), India, for providing research fellowship.

